# Real-time observation of DNA target interrogation and product release by the RNA-guided endonuclease CRISPR Cpf1

**DOI:** 10.1101/205575

**Authors:** Digvijay Singh, John Mallon, Anustup Poddar, Yanbo Wang, Ramreddy Tippana, Olivia Yang, Scott Bailey, Ha Taekjip

## Abstract

CRISPR-Cas9, which imparts adaptive immunity against foreign genomic invaders in certain prokaryotes, has been repurposed for genome-engineering applications. More recently, another RNA-guided CRISPR endonuclease called Cpf1 (also known as Cas12a) was identified and is also being repurposed. Little is known about the kinetics and mechanism of Cpf1 DNA interaction and how sequence mismatches between the DNA target and guide-RNA influence this interaction. We have used single-molecule fluorescence analysis and biochemical assays to characterize DNA interrogation, cleavage, and product release by three Cpf1 orthologues. Our Cpf1 data are consistent with the DNA interrogation mechanism proposed for Cas9, they both bind any DNA in search of PAM (protospacer-adjacent motif) sequences, verifies the target sequence directionally from the PAM-proximal end and rapidly rejects any targets that lack a PAM or that are poorly matched with the guide-RNA. Unlike Cas9, which requires 9 bp for stable binding and ~16 bp for cleavage, Cpf1 requires ~ 17 bp sequence match for both stable binding and cleavage. Unlike Cas9, which does not release the DNA cleavage products, Cpf1 rapidly releases the PAM-distal cleavage product, but not the PAM-proximal product. Solution pH, reducing conditions and 5’ guanine in guide-RNA differentially affected different Cpf1 orthologues. Our findings have important implications on Cpf1-based genome engineering and manipulation applications.

---

In bacteria, CRISPR (clustered regularly interspaced short palindromic repeats)–Cas (CRISPR-associated) acts as an adaptive defense system against foreign genetic elements(1). The system achieves adaptive immunity by storing short sequences of invader DNA into the host genome, which get transcribed and processed into small CRISPR RNA (crRNA). These crRNAs form a complex with a CRISPR nuclease to guide the nuclease to complementary foreign nucleic acids (protospacers) for cleavage. Binding and cleavage also require that the protospacer be adjacent to the protospacer adjacent motif (PAM)(2, 3). CRISPR-Cas9, chiefly the Cas9 from *Streptococcus pyogenes (SpCas9)*, has been repurposed to create an RNA-programmable endonuclease for gene knockout and editing(4-6). Nuclease deficient Cas9 has also been used for tagging genomic sites in wide-ranging applications(4-6). This repurposing has revolutionized biology and sparked a search for other CRISPR-Cas enzymes(7, 8). One such search led to the discovery of the Cas protein Cpf1, with some of its orthologues reporting highly specific cleavage activities in mammalian cells(9-12).

Compared to Cas9, Cpf1 has an AT rich PAM (5’-YTTN-3’ vs. 5’-NGG-3’ for SpCas9), a longer protospacer (24 bp vs. 20 bp for Cas9), creates staggered cuts distal to the PAM vs. blunt cuts proximal to the PAM by Cas9(9) (Fig. 1A), and is an even simpler system than Cas9 because it does not require a trans-activating RNA for nuclease activity or guide-RNA maturation(13). Off-target effects remain one of the top concerns for CRISPR-based applications but Cpf1 is reportedly more specific than Cas9(10, 11). However, its kinetics and mechanism of DNA recognition, rejection, cleavage and product release as a function of mismatches between the guide-RNA and target DNA remain unknown. Precise characterization of differences amongst different CRISPR enzymes should help in expanding the functionalities of the CRISPR toolbox.

**Figure 1.**
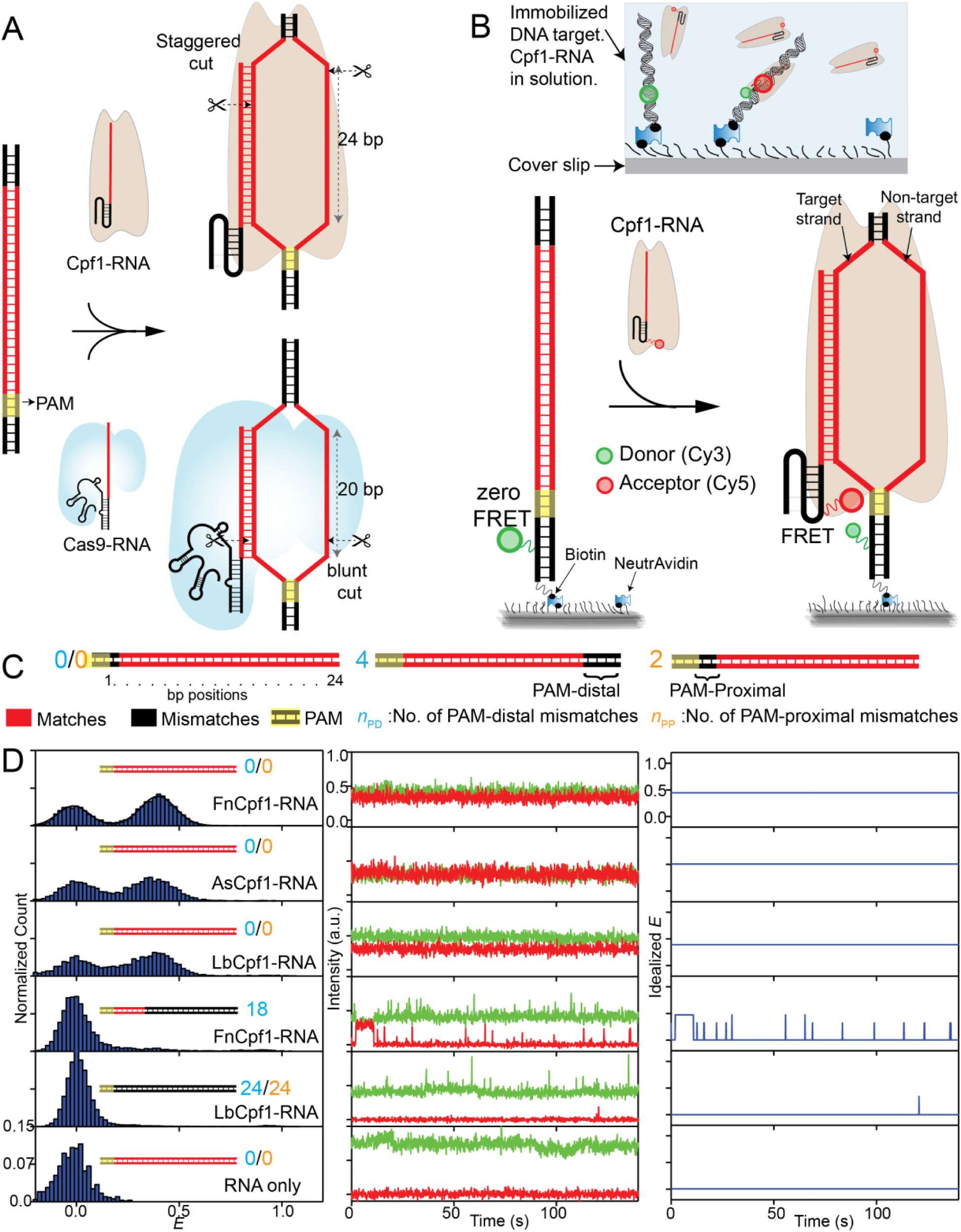
smFRET assay to study DNA interrogation by Cpf1-RNA. (A) Schematic of DNA targeting by CRISPR-Cpf1 and CRISPR-Cas9, and comparison between them. In recent structures (27, 30, 32), the last 4 PAM-distal bp were not unwound and without any RNA-DNA base-pairing for some orthologs. It is unknown whether this is a common feature of all Cpf1 enzymes and currently protospacer for Cpf1 is still taken to be 24 bp long. (B) Schematic of single-molecule FRET assay. Cy3-labeled DNA immobilized on a passivated surface is targeted by a Cy5-labeled guide-RNA in complex with Cpf1, referred to as Cpf1-RNA. (C) DNA targets with mismatches in the protospacer region against the guide-RNA. The number of mismatches PAM-distal (*n*_PD_) and PAM-proximal (*n*_PP_) are shown in cyan and orange, respectively. (D) *E* histograms (left) at 50 nM Cpf1-RNA or 50 nM RNA only. Representative single molecule intensity time traces of donor (green) and acceptor (red) are shown (middle), along with *E* values idealized (right) by hidden Markov modeling(29).

Here, we have used single-molecule fluorescence analysis and biochemical assays to understand how mismatches between the guide-RNA and DNA target modulate the activity of three Cpf1 orthologues from *Acidaminococcus sp*. (AsCpf1), *Lachnospiraceae bacterium* (LbCpf1) and *Francisella novicida* (FnCpf1)(9). Single-molecule methods have been helpful in the study of CRISPR mechanisms(14-23) because they allow real time detection of reaction intermediates and transient states (24).

## Results

### Real time DNA interrogation by Cpf1-RNA

We employed a single-molecule fluorescence resonance energy transfer (smFRET) binding assay(25, 26). DNA targets (donor-labeled, 82 bp long) were immobilized on a polyethylene glycol (PEG) passivated surface and Cpf1 pre-complexed with acceptor-labeled guide-RNA (Cpf1-RNA) was added. Cognate DNA and guide-RNA sequences are identical to the Cpf1 orthologue-specific sequences that were previously characterized biochemically(9) with the exception that we used canonical guide-RNA of AsCpf1 for FnCpf1 analysis because guide-RNAs of AsCpf1 and FnCpf1 are interchangeable(9) (**Supplementary Fig. 1**). Locations of donor (Cy3) and acceptor (Cy5) fluorophores were chosen such that FRET would report on interaction between the DNA target and Cpf1-RNA(27) (Fig. 1B and **Supplementary Fig. 1**). Fluorescent labeling did not affect cleavage activity of Cpf1-RNA (**Supplementary Fig. 2**). We used a series of DNA targets containing different degrees of mismatches relative to the guide-RNA referred to here with *n*_PD_ (the number of PAM-distal mismatches) or *n*_PP_ (the number of PAM-proximal mismatches) (Fig. 1C).

**Figure 2.**
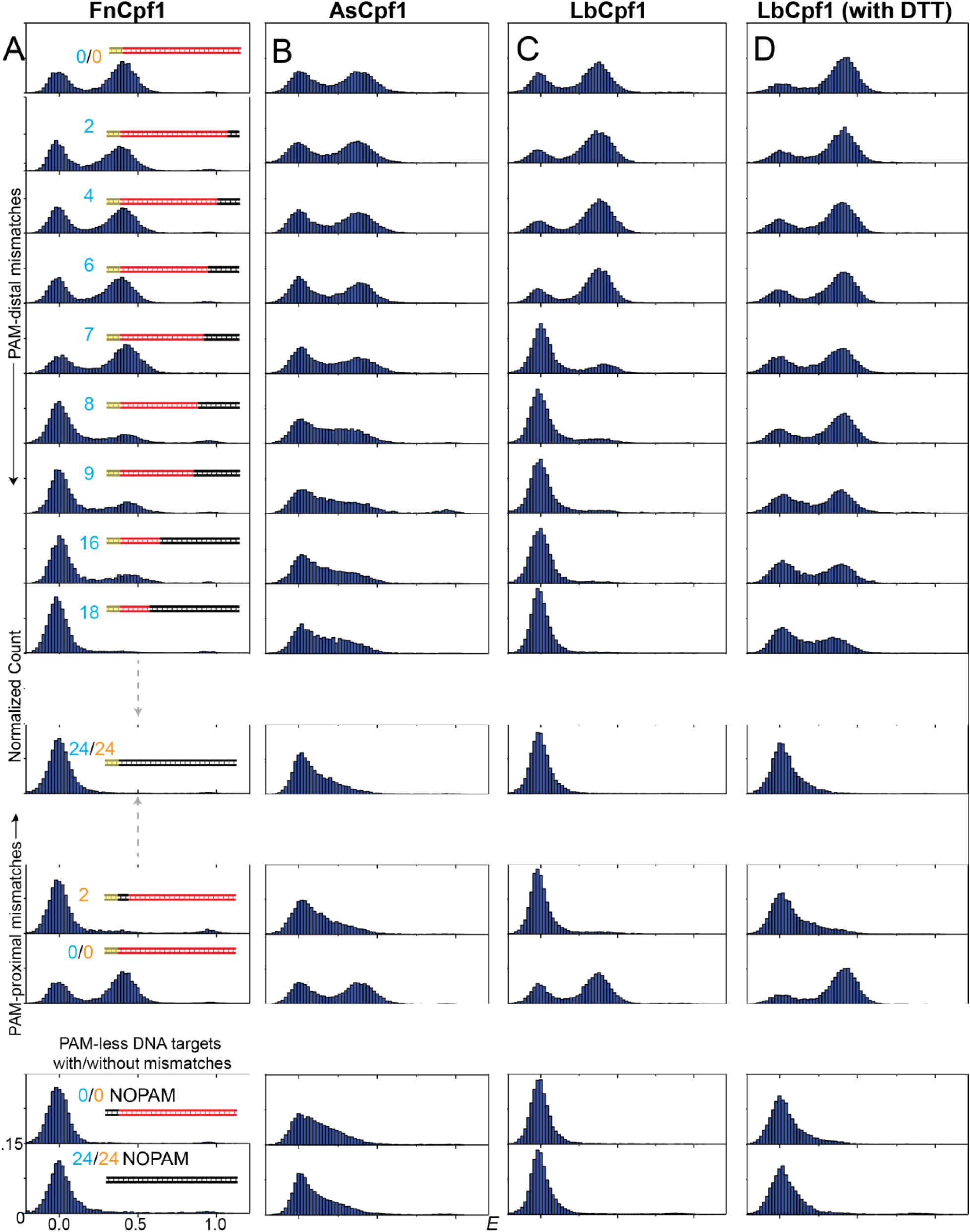
*E* histograms during DNA interrogation by Cpf1-RNA. (A) FnCpf1. (B) AsCpf1. (C) LbCpf1. (D) LbCpf1 (in reducing conditions of 5 mM DTT). Number of PAM-distal (*n*_PD_) and PAM-proximal mismatches (*n*_PP_) are shown in cyan and orange respectively. [Cpf1-RNA] =50 nM. The third peak at high FRET efficiencies occurred only some experiments and was the result of fluorescent impurities likely due variations in PEG passivation, and they were difficult to exclude in automated analysis.

Cognate DNA target in the presence of 50 nM Cpf1-RNA gave two distinct populations with FRET efficiency *E* centered at 0.4 and 0. Using instead a non-cognate DNA target (*n*_PD_ of 24 and without PAM) or guide-RNA only without Cpf1 gave a negligible *E*=0.4 population, allowing us to assign *E*~0.4 to a sequence-specific Cpf1-RNA-DNA complex where the labeling sites are separated by 54 Å(27) (Fig. 1D and **Supplementary Fig. 1**). The *E*=0 population is a combination of unbound states and bound states but with an inactive or missing acceptor. smFRET time trajectories of the cognate DNA target showed a constant *E*~0.4 value within measurement noise (Fig. 1D).

Cpf1-RNA titration experiments yielded dissociation constants (*K*_d_) of 0.27 nM (FnCpf1), 0.1 nM (AsCpf1), 3.9 nM (LbCpf1) in our standard imaging condition and 0.13 nM (LbCpf1) in a reducing condition (**Supplementary Fig. 3**). Binding is much tighter than the 50 nM *K*_d_ previously reported for FnCpf1(13). We performed purification and biochemical experiments in buffer containing dithiothreitol (DTT) as per previous protocols(9) but did not include DTT for standard imaging condition because of severe fluorescence intermittency of Cy5 caused by DTT (28). DTT did not affect FnCpf1 or AsCpf1 DNA binding but made binding >20-fold tighter for LbCpf1 (**Supplementary Fig. 3**). Cleavage by AsCpf1 is most effective at pH 6.5-7.0 (**Supplementary Fig. 4**). Therefore, we used pH 7.0 for AsCpf1 and standard pH 8.0 for FnCpf1 and LbCpf1.

**Figure 3.**
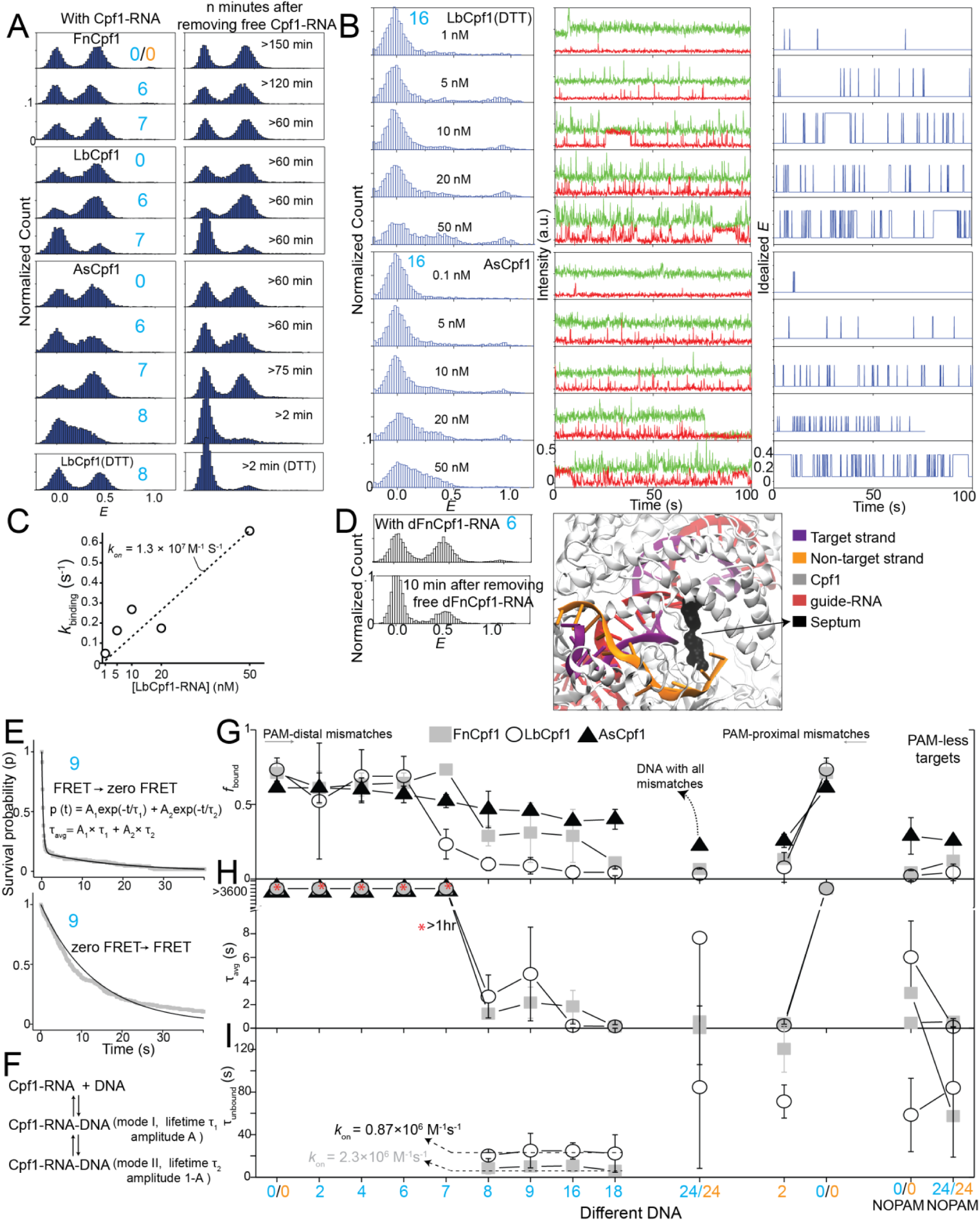
Dynamic interaction of Cpf1-RNA with DNA as a function of mismatches. (A) *E* histograms for various *n*_PD_ with 50 nM Cpf1-RNA (left) and indicated minutes after free Cpf1-RNA was washed out (right) for FnCpf1, LbCpf1, AsCpf1 and LbCpf1 in reducing condition of 5 mM DTT. (B) *E* histograms (left) and representative smFRET time trajectories (middle) with their idealized *E* values (right) for *n*_PD_ = 16 at various concentrations of LbCpf1-RNA in reducing condition and AsCpf1-RNA. The third peak at high FRET efficiencies occurred only some experiments and was the result of fluorescent impurities likely due variations in PEG passivation, and they were difficult to exclude in automated analysis. (C) Rate of LbCpf1-RNA and DNA association (*k*_binding_) at different LbCpf1-RNA concentration. *E* > 0.2 and *E* <0.2 states were taken as putative bound and unbound states. Dwell-times of the unbound states were used to calculate *k*_binding_. (D) Compared to FnCpf1, dFnCpf1 dissociates much quicker from DNA as shown by the change in bound population with and after removal of free dFnCpf1-RNA (left). A septum, preventing the re-hybridization of target and non-target strand, emerges after DNA cleavage which could prevent dissociation of Cpf1-RNA (PDB ID:5MGA(30)). (E) Survival probability of FRET state (*E* > 0.2; putative bound states) and zero FRET state (E<0.2; unbound states) dwell times vs. time, fit with double-exponential and single exponential decay to obtain lifetime of bound state (*τ*_avg_) and unbound state (*τ*_unbound_), respectively. (F) A Model describing a bimodal binding nature of Cpf1-RNA. (G) *f*_bound_, (H) bound state lifetime, (I) unbound state lifetime for various mismatches at 50 nM Cpf1-RNA. Average of rates of binding (*τ*_unbound_^-1^) of DNA with *n*_PD_ = 8-18 were used to calculate *kon* for FnCpf1 and LbCpf1. *n*_*PD*_ and *n*_*PP*_ are shown in cyan and orange, respectively.

**Figure 4.**
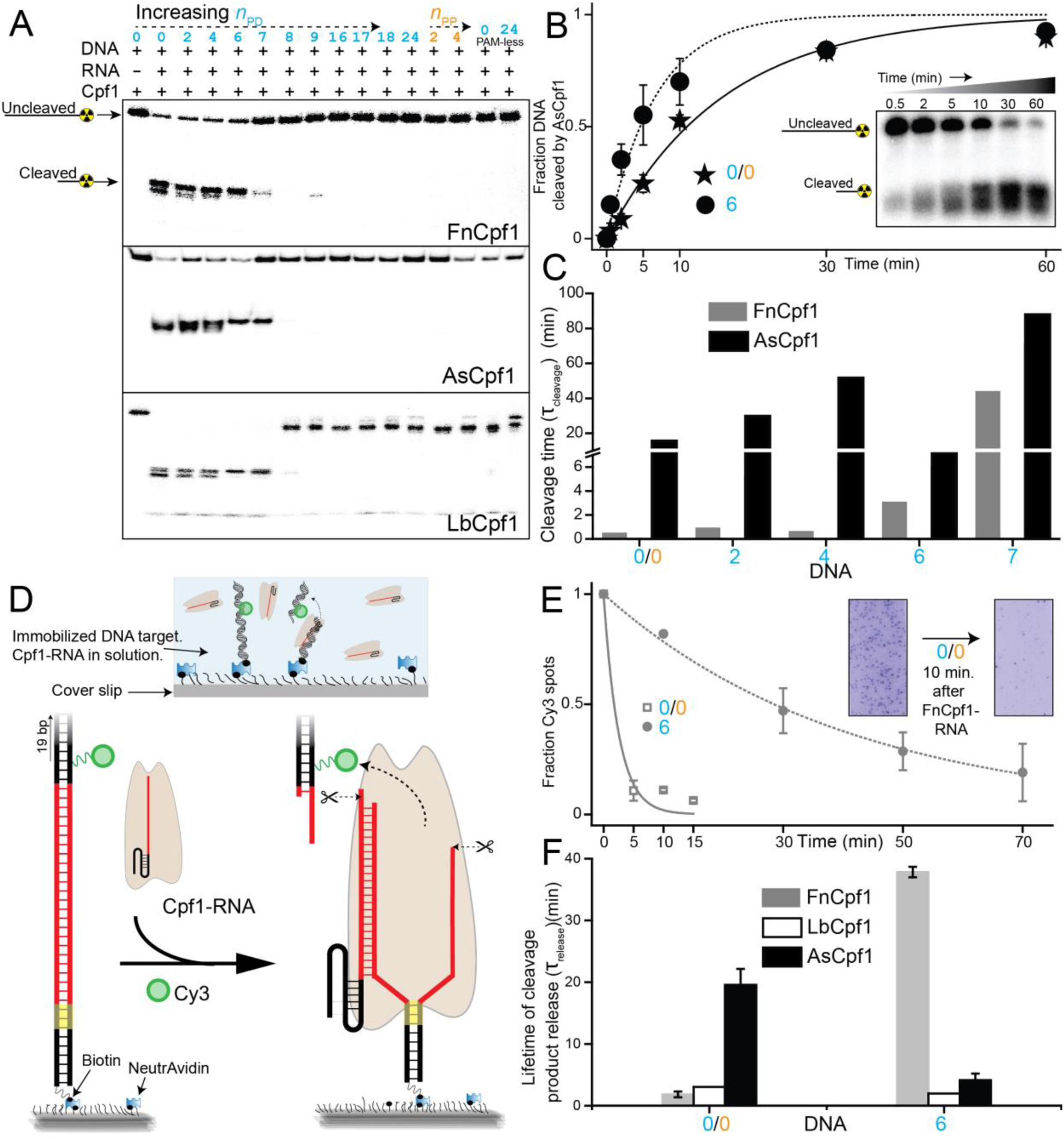
DNA cleavage and product release. (A) Cpf1 induced DNA cleavage at room temperature analyzed by 10% denaturing polyacrylamide gel electrophoresis of radio-labeled DNA targets. (B) Fraction of DNA cleaved by AsCpf1 vs. time for cognate and DNA with *n*_PD_=6, and single exponential fits. A representative gel image is shown in inset. (C) Cleavage time (*τ*_cleavage_) determined from cleavage time courses as shown in (b). (D) Schematic of single-molecule cleavage product release assay. PAM-distal cleavage product release can be detected as disappearance of fluorescence signal from Cy3 attached to the PAM-distal product. (E) Average fraction of Cy3 spots remaining vs time for FnCpf1-RNA (50 nM). Inset shows images before and after 10 min reaction. (F) Average time of cleavage product release (*τ*_release_).

*E* histograms obtained at 50 nM Cpf1-RNA show the impact of mismatches on DNA binding (Fig. 2). The apparent bound fraction *f*_bound_, defined as the fraction of DNA molecules with *E* > 0.2, remained unchanged when *n*_PD_ increased from 0 to 7 (0 to 6 for LbCpf1 in non-reducing conditions) (Fig. 2 and **3G**). Binding was ultrastable for *n*_PD_ ≤ 7; *f*_bound_ did not change even one hour after washing away free Cpf1-RNA (Fig. 3A). *f*_bound_ decreased steeply when *n*_PD_ exceeded 7 for FnCpf1 and LbCpf1 but the decrease was gradual for AsCpf1 and for LbCpf1 in the reducing condition (Fig. 2 and **3G**). For all Cpf1 orthologues, ultrastable binding required *n*_PD_ ≤ 7, corresponding to a 17 bp PAM-proximal sequence match. This is much larger than the 9 bp PAM-proximal sequence match required for ultrastable binding of Cas9(19). PAM-proximal mismatches are highly deleterious for Cpf1 binding because *f*_bound_ dropped by more than 95 % if *n*_PP_ ≥ 2 (Fig. 2 and **3G**). In comparison, Cas9 showed a more modest ~50 % drop for *n*_PP_ = 2 (19). Overall, Cpf1 is much better than Cas9 in discriminating against both PAM-distal and PAM-proximal mismatches for stable binding.

Single molecule time trajectories of all Cpf1 orthologues for *n*_PD_ ≤ 7 showed a constant *E*~0.4 value within noise, limited only by photobleaching. For *n*_PD_ > 7, we observed reversible transitions in *E* likely due to transient binding (**Supplementary Fig. 5-7**). Dwell time analysis as a function of Cpf1-RNA concentration confirmed that *E* fluctuations are due to binding and dissociation, not conformational changes (Fig. 3B, **3C**, and **Supplementary Fig. 3**). We used hidden Markov modeling analysis(29) to segment the time traces to bound and unbound states. Average lifetime of the bound state, *τ*_avg_, was > one hour for *n*_PD_ ≤ 7 but decreased to a few seconds with *n*_PD_ > 7 or any PAM-proximal mismatches (Fig. 3H). The unbound state lifetime differed between orthologues but was nearly the same among most DNA targets, indicating that initial binding has little sequence dependence. The bimolecular association rate *k*_on_ was 2.37 × 10^6^ M^-1^ s^-1^ (FnCpf1), 0.87 × 10^6^ M^-1^ s^-1^ (LbCpf1) and 1.33 × 10^7^ M^-1^ s^-1^ (LbCpf1 in reducing conditions) (Fig. 3C, I). Much longer apparent unbound state lifetimes with PAM-proximal mismatches or DNA targets without PAM are likely due to binding events shorter than the time resolution (0.1s).

**Figure 5.**
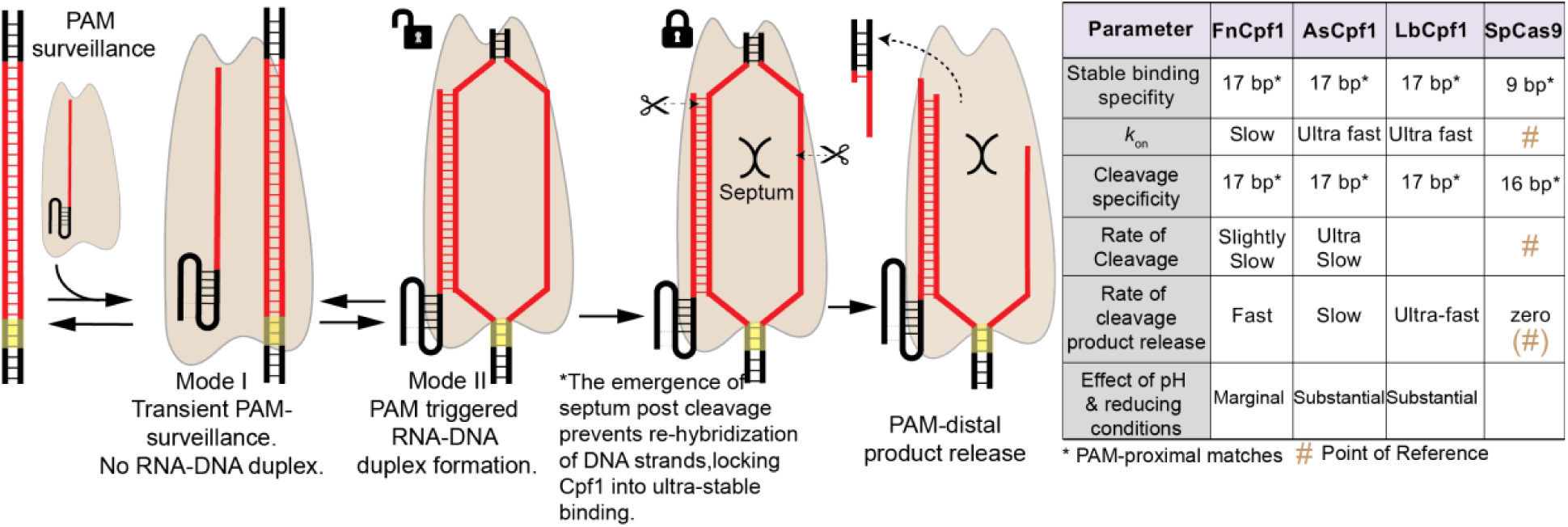
Model of Cpf1-RNA DNA targeting, cleavage and product release.

These results indicate that Cpf1-RNA has dual binding modes. It first binds DNA non-specifically (mode I) in search of PAM and upon detection of PAM, RNA-DNA heteroduplex formation ensues (mode II) and if it extends ≥ 17 bp, Cpf1-RNA remains ultrastably bound to the DNA (Fig. 3F and **Supplementary Fig. 8**). Some reversible transitions in *E* were observed for DNA with *n*_PD_ = 7, indicating that multiple short-lived binding events take place before DNA is cleaved and transitioning to ultrastable binding (**Supplementary Fig. 5-7**and **Supplementary Fig. 13**). RNA-DNA heteroduplex extension is likely directional from PAM-proximal to PAM-distal end because any PAM-proximal mismatch prevented stable binding. Consistent with dual binding modes, survival probability distributions of bound and unbound state were best described by a double and single exponential decay, respectively (Fig. 3E).

### DNA cleavage by Cpf1 as a function of mismatches

Next, we performed gel-based experiments using the same set of DNA targets to measure cleavage by Cpf1. Cleavage was observed at a wide range of temperatures (4-37 °C), required divalent ions (Ca^2+^ could substitute for Mg^2+^), and showed a pH dependence. AsCpf1 is highly active only at slightly acidic to neutral pH (6.5-7.0) whereas FnCpf1 has more activity at pH 8.5 than pH 8.0 (**Supplementary Fig. 9-11**). Cleavage required 17 PAM-proximal matches, corresponding to *n*_PD_ ≤ 7, (Fig. 4A and **Supplementary Fig. 9-10**) which is identical to the threshold for stable binding (Fig. 2 and **3**). This contrasts with Cas9, which requires only 9 PAM-proximal matches for stable binding(19) but 16 PAM-proximal matches for cleavage(3, 14).

We measured the time it takes to cleave DNA, *τ*_cleavage_ (**Supplementary Fig. 12**). *τ*_cleavage_ remained approximately the same among DNA with 0 ≤ *n*_PD_ ≤ 6 for FnCpf1 (30-60 s) but steeply increased upon increasing *n*_PD_ to 7 (Fig. 4B and **4C**). AsCpf1 showed a more complex *n*_PD_ dependence with a minimal *τ*_cleavage_ value of 8 minutes for *n*_PD_ = 6. (Fig. 4C). *τ*_cleavage_ is much longer than the 1 to 15 seconds it takes for Cpf1-RNA to bind the DNA at the same Cpf1-RNA concentration, suggesting that Cpf1-RNA-DNA undergoes additional rate-limiting steps after DNA binding and before cleavage. These additional steps are likely the conformational rearrangement of Cpf1-RNA-DNA complex that position the nuclease domains and DNA strands for cleavage, as has been described in structural analysis of Cpf1-RNA-DNA complex(27, 30).

Because *τ*_cleavage_ is shorter than 60 s for FnCpf1 on DNA targets with *n*_PD_ < 7, we can infer that the ultrastable binding observed for FnCpf1 on the same DNA (lifetime > one hour) is to the cleaved product. Therefore, it is in principle possible that cleavage stabilizes Cpf1-RNA binding and that before cleavage Cpf1-RNA binds to the target DNA less stably. In order to test this possibility, we purified catalytically dead FnCpf1 (dFnCpf1; D917A mutation)(9) and performed the DNA interrogation experiment. dFnCpf1 binding was ultrastable for cognate DNA but showed a substantial dissociation after 5-10 min for *n*_PD_=6 or 7 (Fig. 3D and **Supplementary Fig. 13**). Therefore, cleavage can further stabilize Cpf1-RNA binding to DNA. A septum separating the target and non-target strands and preventing their re-hybridization was observed only after cleavage in Cpf1-RNA-DNA structure (27, 30-32). The formation of this septum during/after DNA cleavage could be the basis of higher stability of Cpf1-RNA binding to DNA post cleavage. Cleavage was negligible for DNA targets that showed transient binding. Therefore, transient binding and dissociation we observed is not to and from a cleaved DNA product.

### Fate of cleaved DNA

For the downstream processing of a cleaved DNA, the cleaved site needs to be exposed(33). To investigate the fate of the target DNA after cleavage, we relocated the Cy3 label to the PAM-distal DNA segment that would depart the imaging surface if the Cpf1 releases the cleavage product(s) (Fig. 4D and **Supplementary Fig. 14**). The number of fluorescent spots decreased over time (Fig. 4E), suggesting the cleavage product is released under physiological conditions, which is in stark contrast to Cas9, which holds onto the cleaved DNA and does not release except in denaturing condition (14, 19). Cpf1 releases only the PAM-distal cleavage product, however, because when Cy3 is attached to a site on the PAM-proximal cleavage product, the number of fluorescence spots did not decrease over time (Fig. 1-3). The average time for fluorescence signal disappearance ranged from ~30 s to 30 min depending on the PAM-distal mismatches and Cpf1 orthologues. By subtracting the time it takes to bind and cleave, we estimated the product release time scale (*τ*_release_) (Fig. 4F), which showed a dependence on *n*_PD_. Therefore, PAM-distal mismatches can also affect product release.

## Discussion

The two-step mechanism of sampling for PAM followed by directional RNA-DNA heteroduplex extension (Fig. 5) is shared between Cas9 and Cpf1, suggesting this to be a general target identification mechanism of these CRISPR systems. Ultrastable binding of Cpf1 requires the same extent of sequence match (17 bp PAM-proximal matches) as target cleavage. This contrasts with Cas9, which requires only bp and 16 bp PAM-proximal matches for ultrastable binding and cleavage, respectively(19, 34, 35). Therefore, Cpf1 can be more sequence specific in experiments involving the use of catalytically dead CRISPR enzymes for imaging, tracking and transcription regulation purposes(36). The binding specificity of engineered Cas9s (eCas9(37) & Cas9-HF1(38)) is still much lower than that of Cpf1(35). Therefore, Cpf1 has the potential to be a better alternative to all current Cas9 variants.

Cleavage rate is reduced with increasing PAM-distal mismatches (Fig. 4C) even when the mismatches do not affect stable binding (Fig. 3), suggesting that shorter RNA-DNA heteroduplexes result in slower conformational changes required for cleavage activation. Previous studies on Cas9 revealed that mismatches alter the kinetics of DNA unwinding, RNA-DNA heteroduplex extension, and nuclease and proof-reading domain movements(20, 22, 34, 35).

For cognate DNA target, RNA-DNA heteroduplex extension would require unwinding of the parental DNA duplex. We performed cleavage experiments using DNA with PAM-distal mismatched region pre-unwound in order to test the relative importance of parental DNA duplex unwinding and annealing with RNA in cleavage activation. Cpf1 needed much fewer PAM-proximal matches to cleave if the mismatched region is pre-unwound (**Supplementary Fig. 15**) indicating indeed DNA unwinding is likely more important than RNA-DNA heteroduplex in activating cleavage. Accordingly, ssDNA can also be cleaved by Cpf1 (**Supplementary Fig. 15**). Therefore, the role of RNA may primarily be in keeping the DNA unwound through annealing with the target strand.

CRISPR enzymes bend DNA to cause a local kink near the PAM, which acts as a seed for unwinding and heteroduplex extension(27, 39, 40). Perturbing DNA rigidity by introducing a nick near the PAM slowed down cleavage, underscoring the importance of Cpf1-induced DNA bending for cleavage (**Supplementary Fig. 16**). Cas9 causes a larger DNA bend than Cpf1(27, 39), possibly contributing to it higher tolerance of PAM-proximal mismatches in binding and cleavage activity.

Shorter and simpler guide-RNA(9) for Cpf1 could potentially be deleterious for its engineering or extension, as is done for Cas9’s guide-RNA(41). For example, an extra 5’ guanine in the guide-RNA was extremely deleterious for cleavage by LbCpf1 (**Supplementary Fig. 17**), potentially posing problems for applications where guide-RNAs are transcribed using U6/T7 RNA polymerase systems that require first nucleotide in transcribed RNA to be the guanine(42, 43). This problem may be solved by transcribing RNAs with 5’ G containing CRISPR repeat which will be processed out by Cpf1 itself to produce mature guide-RNAs(13) (**Supplementary Fig. 17**).

Cas9 has provided a highly efficient and versatile platform for DNA targeting, but the efficiency of gene knock-in is low(44). Amongst the possible reasons is the inability of Cas9 to release and expose cleaved DNA ends. In contrast, the ability of Cpf1 to release a cleavage product readily, combined with staggered cuts it generates, could in principle increase the knock-in efficiency. Although it remains to be seen how this property affects the downstream processing *in vivo*, we can also envision a scenario where product release by Cpf1 can be detrimental to genome engineering applications. Applying positive twist to the DNA in a Cas9-RNA-DNA complex can release Cas9-RNA from DNA by promoting rewinding of parental DNA duplex(15). Positive supercoiling is generated ahead of a transcribing RNA polymerase(45) and Cas9 holding onto the double strand break product may help build the torsional strain required to eject Cas9-RNA. If the PAM-distal cleavage product is released prematurely as in the case of Cpf1, transcription-induced positive supercoiling cannot build up and the Cpf1-RNA would remain bound stably to the PAM-proximal cleavage product, hiding the cleaved end and preventing efficient knock-in.

High specificity of adaptive immunity by Cpf1 against hypervariable genetic invaders is a little paradoxical. But Cpf1 and Cas9 systems co-exist in many species and thus they likely provide immunity suited to their features, effectively broadening the scope of immunity. Overall, our results establish major different and common features between Cpf1 and Cas9 which can be useful for the broadening of genome engineering applications as well.

## Author contributions

D.S., T.H. designed the experiments. D.S performed single molecule experiments. J.M. performed radio-labeled gel electrophoresis experiments. D.S. performed gel electrophoresis experiments involving SYBR staining of nucleic acids. D.S., J.M., R.T. expressed and purified Cpf1. D.S. prepared DNA and RNA substrates. D.S. wrote the MATLAB package for data analysis and performed it with help from A.P., Y.W. A.P. assisted with the PEG passivation of some slides. O.Y., Y.W. assisted with some experiments. D.S, T.H., S.B. discussed the data. D.S., T.H. wrote the manuscript.

Authors declare no competing financial interests. Correspondence: T.H.

## Acknowledgements

The project was supported by grants from National Science Foundation (PHY-1430124 to T.H.) and National Institutes of Health (GM065367; GM112659 to T.H and GM097330 to S.B.); T.H. is an investigator with the Howard Hughes Medical Institute. J.M is supported by the Nation Institutes of Health Chemical Biology Interface training program (T32GM080189).

## Materials and methods

### DNA targets for smFRET analysis of DNA interrogation

Single-stranded DNA (ssDNA) oligonucleotides were purchased from Integrated DNA Technologies. ssDNA target and non-target (labeled with Cy3) strands and a biotinylated adaptor strand were mixed. The non-target strand was created by ligating two component strands, one with Cy3 and the other containing the protospacer region to avoid having to synthesize modified oligos for each mismatch construct. For schematics, see **Supplementary Fig. 1A.** Fully duplexed DNA targets but with a nick were also used. The Cy3 fluorophore is located 4 bp upstream of the protospacer adjacent motif (PAM: 5’-YTTN-3’) and was conjugated via Cy3 N-hydroxysuccinimido (Cy3-NHS; GE Healthcare) to the Cy3 oligo at amino-group attached to a modified thymine through a C6 linker (amino-dT) using NHS ester linkage. smFRET experiments were done with both sets of DNA targets (with or without a nick) and no significant differences were found between them. **Supplementary Table 1** shows all DNA targets used. Additional details about the DNA targets is available in the supplementary document.

### DNA targets for real time single-molecule assay for interrogating fate of cleaved DNA

For single-molecule cleavage product release experiments, a non-target strand with the Cy3 relocated in a different position was used. Cy3 label was conjugated onto the amine modification (amino-dT) using Cy3-NHS, as described above. Schematic of these DNA targets is in the **Supplementary Fig. 14** and their sequences in **Supplementary Table 5**.

### DNA targets for gel electrophoresis experiments

They were prepared and hybridized as described above. For radio-labeled gel electrophoresis experiments, the target strand was 5′ radiolabeled with T4 polynucleotide kinase (New England BioLabs) and γ-^32^P ATP (Perkin Elmer). The target and non-target strands were annealed with the non-target strands in excess.

### Guide-RNA

For single molecule experiments, guide-RNA was purchased from IDT with modifications for Cy5 labeling as described in **Supplementary Table 5.** Cy5 was conjugated via Cy5 N-hydroxysuccinimido (Cy5-NHS; GE Healthcare) to the RNA as described previously(19, 46). For all other experiments, unmodified guide-RNA was used and they were either *in vitro* transcribed or purchased from IDT. Guide-RNA sequences used in this study is available in **Supplementary table 5**.

### Preparation of Cpf1-RNA

The Cpf1-RNA was freshly prepared prior to each experiment by mixing the guide-RNA (50 nM) and Cpf1 in 1:3.5 ratio in the following reaction buffers and incubated for at least 10 min at room temperature. 50 mM Tris-HCl (pH 8.0) 100 mM NaCl, 10 mM MgCl_2_, (FnCpf1 and LbCpf1) and 50 mM HEPES (pH 7.0) 100 mM NaCl, 10 mM MgCl_2_, (AsCpf1). 5mM DTT was only used in the buffer when specified. 0.2 mg/ml Bovine serum albumin (BSA), 1 mg/ml glucose oxidase, 0.04 mg/ml catalase, 0.8% dextrose and saturated Trolox (>5 mM) were additional contents of the reaction buffers for single-molecule fluorescence experiments. Excess Cpf1 was used to achieve highest extent of complexation of all the available guide-RNA and the concentration of guide-RNA was used as the concentration of Cpf1-RNA. Cpf1 activity using the similar guide-RNA and on DNA targets with same protospacer and PAM have been characterized previously(9). Fluorophore labeling of either DNA targets or guide-RNA did not impair Cpf1 activity. (**Supplementary Fig. 2**).

### Expression-purification of Cpf1 and Single-molecule detection

These methods have been described previously(9, 26). Their full details are available in the supplementary document.

### FRET efficiency histograms and Cpf1-RNA bound DNA fraction

A smFRET time-trajectory is a series of *E* values every 100 ms. First five *E* values of each single-molecule trace were pooled together to build single molecule *E* histograms. Cpf1-RNA bound DNA fraction (*f*_bound_) was calculated as a ratio between the number of molecules with *E* > 0.2 and the total number of molecules in the *E* histograms. *E* histograms shown in Fig. 2 were constructed by combining data from two independent experiments (except for AsCpf1; PAM-less DNA). At least 2000 molecules, in most cases > 4000, were used for each histogram. The criteria for the selection of fluorescent single molecule spot was same as described previously. Majority of selected spots (~85 %), were used for the analysis. The remaining (~15 %) were discarded as their intensities were too low (likely due to impurities) or too high (impurities, aggregates or multiple fluorescent molecules in a single spot).

### Determination of binding kinetics

For DNA targets that showed real-time reversible binding/dissociation of Cpf1-RNA, idealization of smFRET traces via hidden Markov Model(29) analysis yielded two pre-dominant FRET states, of zero (*E*< 0.2) and bound state (*E*> 0.2). Lifetime of the unbound state, *τ*_unbound_, was calculated by fitting survival probability of dwell times of unwound state (*E*< 0.2) vs time to a single-exponential decay (exp[-t/*τ*_unbound_]). The survival probability of the bound state required a double-exponential decay for adequate fitting (A*exp[-t/*τ*_1_]+ [1-A] *exp[-t/*τ*_2_], and the average lifetime was calculated as *τ*_avg_ = A*τ*_1_ +(1-A)*τ*_2_. At least 60 long-lived smFRET traces, in most cases > 90, were used for the indicated lifetime analysis(s). The bimolecular association rate constant *k*_on_, binding rate *k*_binding_ and dissociation rate *k*_off_ were calculated as follows.

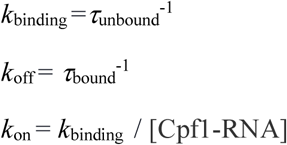

Due to under-sampled binding events, *τ*_avg_ of FnCpf1 for PAM-less DNA and DNA with 2 *n*_PP_ were calculated as the algebraic average of *E*> 0.2 dwell-times. Cy5 labeling efficiency of guide-RNA was ~90% and thus *f*_bound_ and*τ*_unbound_ were appropriately corrected. Due to high noise, the smFRET traces from experiments involving AsCpf1 could not be idealized with high accuracy thus preventing their *k*_off_ and *k*_on_ analysis.

### Estimation of dissociation constant (*Kd*)

To estimate *K*_d_, Cpf1-RNA bound DNA fraction (*f*_bound_) vs Cpf1-RNA concentration (*c*) was fit using *f*_bound_= M × c / (*K_d_* + c) where M is the maximum observable *f*_bound_. M is typically less than 1 because inactive or missing acceptors or because not all of the DNA on the surface are capable of binding Cpf1-RNA.

### Overall lifetime of release of cleavage products

Single-molecule experiments were used to estimate the lifetime of the release of cleavage products by fitting the decreasing number of Cy3 spots (loss of spots due to Cpf1-RNA induced cleavage and release) to a single-exponential decay. The time of binding (*k*_on_×50 nM) and time of cleavage (*τ*_cleavage_) were subtracted from the obtained lifetime to get the true lifetime of the release (*τ*_release_) of cleavage products. But since *τ*_cleavage_ was not measured for LbCpf1, its reported *τ*_release_ is without the *τ*_cleavage_ and time of binding subtraction.

### Gel electrophoresis experiments involving visualization of nucleic acid bands via SYBR Gold II staining

All experiments were conducted by mixing DNA targets and Cpf1-RNA in 1:5 ratio in the reaction buffer. The reaction was incubated for 4.5-5 hr (unless stated otherwise) before being resolved by 4% native/denaturing agarose gel electrophoresis and SYBR Gold II staining of nucleic acids using the precast gels containing SYBR Gold II, purchased from Thermo Fisher Scientific. For native gel electrophoresis, the reaction aliquots were directly loaded onto the gels. All the reactions were incubated at the room temperature, 37 °C or 4 °C and indicated in the presentation of their results. The gel electrophoresis was run at room temperature for experiments incubated at room temperature/37 °C and at 4 °C for experiments incubated at 4 °C. The cleaved-uncleaved DNA target with/without the bound Cpf1-RNA along with other nucleic acids were stained by SYBR Gold II and imaged by blue laser illumination (480 nm; GE Amersham Molecular Dynamics Typhoon 9410 Molecular Imager and 488 nm; Amersham Imager 600). For all of these experiments, the concentration of the DNA targets ranged from 20 nM to 60 nM and consequently the effective concentration of Cpf1-RNA ranged from 100 nM to 300 nM respectively. Volume of aliquots used for gel loading ranged from 10 to 20 μL per lane. For the time-lapse denaturing gel electrophoresis experiments, the acquired gel-images were quantified using ImageJ(47). Entire panel of DNA targets used in these gel-electrophoresis experiments is available in **Supplementary Table 3** and **4**. Tris-HCl at pH 8.0 was used in the reaction buffers for all the experiments except for the ones reported in **Supplementary Fig. 2** and **Supplementary Fig. 9** where Tris-HCl at pH 8.5 was used.

### Gel electrophoresis experiments and autoradiography

Experiments containing radiolabeled DNA substrates were performed as above. However, samples were quenched, in buffer containing 95% formamide, 0.01% SDS 0.01% bromophenol blue, 0.01% xylene cyanol, and 1 mM EDTA and incubated at 95 °C for 5min then on ice for 2min. Volume ratio of quenching buffer to reaction was 5:1. Samples were loaded on to denaturing polyacrylamide gels (10% acrylamide, 50%(w/v) urea) and allowed to separate. Amount of sample loaded on to gel was normalized to 10,000 counts per sample. Gels were imaged via phosphor screens. Entire panel of DNA targets used in these gel-electrophoresis experiments is available in **Supplementary Table 2**. All the gel electrophoresis experiments were done in the following reaction buffers: 50 mM Tris-HCl (pH 8.0) 100 mM NaCl, 10 mM MgCl_2_, 5 mM DTT (FnCpf1 and LbCpf1) and 50 mM HEPES (pH 7.0) 100 mM NaCl, 10 mM MgCl_2_, 5 mM DTT (AsCpf1). For all experiments (single molecule fluorescence analysis and gel electrophoresis experiments), errors bars represent standard deviation from the analysis of 2 or 3 replicate experiments.

## References

1. Marraffini LA & Sontheimer EJ (2010) CRISPR interference: RNA-directed adaptive immunity in bacteria and archaea. Nat Rev Genet 11(3):181–190.

2. Gasiunas G, Barrangou R, Horvath P, & Siksnys V (2012) Cas9-crRNA ribonucleoprotein complex mediates specific DNA cleavage for adaptive immunity in bacteria. Proc Natl Acad Sci U S A 109(39):E2579–2586.

3. Jinek M, et al. (2012) A programmable dual-RNA-guided DNA endonuclease in adaptive bacterial immunity. Science 337(6096):816–821.

4. Barrangou R & Doudna JA (2016) Applications of CRISPR technologies in research and beyond. Nat Biotechnol 34(9):933–941.

5. Wang H, La Russa M, & Qi LS (2016) CRISPR/Cas9 in Genome Editing and Beyond. Annu Rev Biochem 85:227–264.

6. Wright AV, Nuñez JK, & Doudna JA (2016) Biology and Applications of CRISPR Systems: Harnessing Nature’s Toolbox for Genome Engineering. Cell 164(1–2):29–44.

7. Shmakov S, et al. (2017) Diversity and evolution of class 2 CRISPR-Cas systems. Nat Rev Microbiol 15(3):169–182.

8. Burstein D, et al. (2017) New CRISPR-Cas systems from uncultivated microbes. Nature 542(7640):237–241.

9. Zetsche B, et al. (2015) Cpf1 is a single RNA-guided endonuclease of a class 2 CRISPR-Cas system. Cell 163(3):759–771.

10. Kleinstiver BP, et al. (2016) Genome-wide specificities of CRISPR-Cas Cpf1 nucleases in human cells. Nat Biotechnol 34(8):869–874.

11. Kim D, et al. (2016) Genome-wide analysis reveals specificities of Cpf1 endonucleases in human cells. Nat Biotechnol 34(8):863–868.

12. Makarova KS, et al. (2015) An updated evolutionary classification of CRISPR-Cas systems. Nat Rev Microbiol 13(11):722–736.

13. Fonfara I, Richter H, Bratovič M, Le Rhun A, & Charpentier E (2016) The CRISPR-associated DNA-cleaving enzyme Cpf1 also processes precursor CRISPR RNA. Nature 532(7600):517–521.

14. Sternberg SH, Redding S, Jinek M, Greene EC, & Doudna JA (2014) DNA interrogation by the CRISPR RNA-guided endonuclease Cas9. Nature 507(7490):62–67.

15. Szczelkun MD, et al. (2014) Direct observation of R-loop formation by single RNA-guided Cas9 and Cascade effector complexes. Proc Natl Acad Sci U S A 111(27):9798–9803.

16. Redding S, et al. (2015) Surveillance and Processing of Foreign DNA by the Escherichia coli CRISPR-Cas System. Cell 163(4):854–865.

17. Rutkauskas M, et al. (2015) Directional R-Loop Formation by the CRISPR-Cas Surveillance Complex Cascade Provides Efficient Off-Target Site Rejection. Cell Rep.

18. Josephs EA, et al. (2016) Structure and specificity of the RNA-guided endonuclease Cas9 during DNA interrogation, target binding and cleavage. Nucleic Acids Res 44(5):2474.

19. Singh D, Sternberg SH, Fei J, Doudna JA, & Ha T (2016) Real-time observation of DNA recognition and rejection by the RNA-guided endonuclease Cas9. Nat Commun 7:12778.

20. Lim Y, et al. (2016) Structural roles of guide RNAs in the nuclease activity of Cas9 endonuclease. Nat Commun 7:13350.

21. Blosser TR, et al. (2015) Two distinct DNA binding modes guide dual roles of a CRISPR-Cas protein complex. Mol Cell 58(1):60–70.

22. Dagdas YS, Chen JS, Sternberg SH, Doudna JA, & Yildiz A (2017) A Conformational Checkpoint Between DNA Binding And Cleavage By CRISPR-Cas9. bioRxiv.

23. Chen JS, et al. (2017) Enhanced proofreading governs CRISPR-Cas9 targeting accuracy. bioRxiv.

24. Joo C, Balci H, Ishitsuka Y, Buranachai C, & Ha T (2008) Advances in single-molecule fluorescence methods for molecular biology. Annu Rev Biochem 77:51–76.

25. Ha T, et al. (1996) Probing the interaction between two single molecules: fluorescence resonance energy transfer between a single donor and a single acceptor. Proc Natl Acad Sci U S A 93(13):6264–6268.

26. Roy R, Hohng S, & Ha T (2008) A practical guide to single-molecule FRET. Nat Methods 5(6):507–516.

27. Yamano T, et al. (2016) Crystal Structure of Cpf1 in Complex with Guide RNA and Target DNA. Cell 165:949–962.

28. Bates M, Blosser TR, & Zhuang X (2005) Short-range spectroscopic ruler based on a single-molecule optical switch. Phys Rev Lett 94(10):108101.

29. McKinney SA, Joo C, & Ha T (2006) Analysis of single-molecule FRET trajectories using hidden Markov modeling. Biophys J 91(5):1941–1951.

30. Stella S, Alcon P, & Montoya G (2017) Structure of the Cpf1 endonuclease R-loop complex after target DNA cleavage. Nature 546(7659):559–563.

31. Stella S, Alcon P, & Montoya G (2017) Class 2 CRISPR-Cas RNA-guided endonucleases: Swiss Army knives of genome editing. Nat Struct Mol Biol 24(11):882–892.

32. Swarts DC, van der Oost J, & Jinek M (2017) Structural Basis for Guide RNA Processing and Seed-Dependent DNA Targeting by CRISPR-Cas12a. Mol Cell 66(2):221–233.e224.

33. Richardson CD, Ray GJ, DeWitt MA, Curie GL, & Corn JE (2016) Enhancing homology-directed genome editing by catalytically active and inactive CRISPR-Cas9 using asymmetric donor DNA. Nat Biotechnol 34(3):339–344.

34. Sternberg SH, LaFrance B, Kaplan M, & Doudna JA (2015) Conformational control of DNA target cleavage by CRISPR-Cas9. Nature 527(7576):110–113.

35. Singh D, et al. (2017) Mechanisms of improved specificity of engineered Cas9s revealed by single molecule analysis. bioRxiv.

36. Doudna JA & Charpentier E (2014) Genome editing. The new frontier of genome engineering with CRISPR-Cas9. Science 346(6213):1258096.

37. Slaymaker IM, et al. (2016) Rationally engineered Cas9 nucleases with improved specificity. Science 351(6268):84–88.

38. Kleinstiver BP, et al. (2016) High-fidelity CRISPR-Cas9 nucleases with no detectable genome-wide off-target effects. Nature 529(7587):490–495.

39. Jiang F, et al. (2016) Structures of a CRISPR-Cas9 R-loop complex primed for DNA cleavage. Science 351(6275):867–871.

40. Anders C (2014) Structural basis of PAM-dependent target DNA recognition by the Cas9 endonuclease. 513(7519):569–573.

41. Wang S, Su JH, Zhang F, & Zhuang X (2016) An RNA-aptamer-based two-color CRISPR labeling system. Sci Rep 6:26857.

42. Oakley JL & Coleman JE (1977) Structure of a promoter for T7 RNA polymerase. Proc Natl Acad Sci U S A 74(10):4266–4270.

43. Guschin DY, et al. (2010) A rapid and general assay for monitoring endogenous gene modification. Methods Mol Biol 649:247–256.

44. Kan Y, Ruis B, Lin S, & Hendrickson EA (2014) The mechanism of gene targeting in human somatic cells. PLoS Genet 10(4):e1004251.

45. Revyakin A, Liu C, Ebright RH, & Strick TR (2006) Abortive initiation and productive initiation by RNA polymerase involve DNA scrunching. Science 314(5802):1139–1143.

46. Joo C & Ha T (2012) Labeling DNA (or RNA) for single-molecule FRET. Cold Spring Harb Protoc 2012(9):1005–1008.

47. Schneider CA, Rasband WS, & Eliceiri KW (2012) NIH Image to ImageJ: 25 years of image analysis. Nat Methods 9(7):671–675.

